# Deciphering the molecular mechanism of the cancer formation by chromosome structural dynamics

**DOI:** 10.1101/2021.02.15.431330

**Authors:** Xiakun Chu, Jin Wang

## Abstract

Cancer reflects the dysregulation of the underlying gene network, which is intimately related to the 3D genome organization. Numerous efforts have been spent on experimental characterizations of the structural alterations in cancer genomes. However, there is still a lack of genomic structural-level understanding of the temporal dynamics for cancer initiation and progression. Here, we use a landscape-switching model to investigate the chromosomal structural transition during the can-cerization and reversion processes. We find that the chromosome undergoes a non-monotonic structural shape-changing pathway with initial expansion followed by compaction during both of these processes. Furthermore, our analysis reveals that the chromosome with a more expanded structure than those at both the normal and cancer cell during cancerization exhibits a sparse contact pattern, which shows significant structural similarity to the one at the embryonic stem cell in many aspects, including the trend of contact probability declining with the genomic distance, the global structural shape geometry and the spatial distribution of loci on chromosome. We show that cell cancerization and reversion are highly irreversible processes in terms of the chromosomal structural transition pathways, spatial repositioning of chromosomal loci and hysteresis loop of contact evolution analysis. Our model draws a molecular-scale picture of cell cancerization, which contains initial reprogramming towards the stem cell followed by differentiation towards the cancer cell, accompanied by an initial increase and subsequent decrease of cell stemness.

## 1 INTRODUCTION

Currently, our knowledge of how cancer cells form and proliferate is still quite limited. In general, the development of cancer is controlled by the underlying gene regulatory network within a cell and cell-cell interactions [1], which rely on the molecular interactions between the spatially organized genome and a broad class of proteins, including the transcription factors and chromatin remodelers [2]. As the structural scaffold for the genome function, the 3D genome architecture has been recognized to play an important role in regulating the gene expression [3,4, 5], leading to the intimate structure-function relationships at the genomic level. Several studies have shown that alterations in chromosome structures and interactions can make significant contributions to the dysregulation of the gene expressions forming the specific cancer signatures [6, 7, 8, 9]. These findings have fostered a view that the 3D context of the genome is the major player in the development and progression of cancer [10]. Therefore, investigating the disorganization of the 3D genome structure in cancer cells can provide the key to understanding the pathogenesis of cancer.

Although the chromosomal structural variant, which is the major form of the genome instability, has been regarded as a hallmark of almost all human cancers [11, 12, 13], determining the genome structure has been a long-term challenge until the emergence of chromosome conformation capture (3C) techniques nearly two decades ago [14]. As an advanced derivative of the 3C, Hi-C measures the spatial proximity of the chromosomal loci across the entire genome in terms of the contact frequency map and provides an ensemble description of the genome organization within a large number of cells [15, 16]. Recently, Hi-C techniques were applied to characterize the disorganization of the cancer genomes and determine their functional consequences [17]. Taberlay et al. observed the smaller-sized topologically associated domains (TADs) formed in prostate cancer cells than those in normal cells due to the emergence of additional domain boundaries. This further leads to alterations of TP53 tumor suppressor locus [18]. In addition, the effects of the structural disruptions within TADs on leading to various cancer types were established [19, 20, 21]. At the higher hierarchical level, Barutcu et al. found that the significant A/B compartment switching between the normal and breast cancer cell is associated with gene expression changes [22]. These studies have shown that the structural variants in the cancer genome occur throughout the genome sequence, and they have important impacts on the gene expression alterations for inducing cancer formation.

Increasing evidence indicates that cancer development is orchestrated by a small subpopulation of the cancer cells, namely the cancer stem (CS) cells. [23, 24, 25, 26]. CS cells, which possess stem-like properties and functions, are capable of performing self-renewal, proliferation and differentiation. Thus they provide the driving forces for cancer progression [27, 28]. However, experimental identifications on the CS cell and the associated dynamics are non-trivial due to its rare population in the cancer cells [29]. The previous theoretical studies using gene regulatory work, which includes the cancer and development genetic markers as well as the interactions between them, provided a landscape view of how the cancer cell develops and CS cell forms [30, 31]. However, the work focused on simplified core gene regulatory networks to describe the cancer systems, so the results were restricted in the form of regulation pathways among several marker genes and this description is at the gene network level. Therefore, there is still a lack of a complete molecular chromosomal-structural level understanding of cancer development and CS cell formation. On the other hand, the strategies on inducing the cancer cell to normal cell by reversion [32] or transdifferentiation processes [33, 34, 35] are rapidly developing. These approaches have opened new avenues for cancer treatments, but the underlying mechanism of the cancer reversion is still unclear [36].

Despite the significant achievements made by the Hi-C technique on elucidating the spatial genome disorganization in the cancer cells [17], the temporal dynamic rearrangement of the genome structure during cancer development is still not available. The picture is fundamentally important as it may provide a molecular-level understanding of the cancer mechanism. However, measuring the chromosomal structural dynamics during the cancer cell developmental process is extremely challenging due to the spatial and temporal resolution limits of the current experiments. Here, we aim to address this issue with the computational efforts by investigating the dynamic cancer-related chromosomal structural evolution, which provides the microscopic description of the cancer process.

We developed molecular dynamics models and performed associated simulations to undertake the task. First, we applied the maximum entropy principle to incorporate the Hi-C data of the normal and cancer cells into two independent sets of simulations. These simulations generated two potentials for describing the chromosome structural ensembles in the normal and cancer cells, respectively. In the cell nucleus, the chromosome constantly experiences the non-equilibrium effects even at one cell state, leading to a non-equilibrium system. Theoretically, it has been demonstrated that the dynamics of the non-equilibrium system can be described using the concept of an effective landscape in some circumstances [37, 38]. Recent studies revealed that under an effective equilibrium landscape, the chromosome dynamics reproduces many aspects of kinetic behaviors as observed in experiments, including anomalous diffusion, viscoelasticity, and spatially coherent dynamics [39, 40]. Here, to examine whether the potentials accounting for chromosome structures obtained from maximum entropy principle simulations can also be used for describing the chromosome dynamics, we performed additional simulations under these two potentials and calculated the diffusion behaviors of the chromosome motion in the normal and cancer cell, respectively (Figure S3). We observed sub-diffusivity of chromosome dynamics with the scaling exponents of the mean square displacement in good agreement with multiple experiments [41, 42]. Our results have suggested that these two potentials generated by the maximum entropy principle simulations for chromosomes in the normal and cancer cell can be further regarded as the effective energy landscapes that govern the chromosome conformational dynamics within the normal and cancer cell, respectively. Although each state can be described by an effective equilibrium landscape, the inter-basin dynamics of switching between normal and cancer states often requires a significant amount of energy input [43] and therefore is a non-equilibrium process, which cannot be treated in an equilibrium way. This is the motivation and target of our current study. Therefore, the cancer-related chromosome structural transitions are further described by the connection between these two effective landscapes during the cell cancerization and reversion processes.

To bridge these two landscapes for describing transitions between the normal and cancer cell, we then used a landscape-switching model, which was developed to simulate the chromosomal structural transition during the cell cycle [44] and the cell developmental processes [45]. The model employed an instantaneous energy excitation to trigger the transition processes followed by the relaxation dynamics (see ‘‘**Materials and Methods**”). The switch implementation in the model has a practical correspondence to mimic the roles of the genetic mutations and epigenetic modifications in initiating cancer processes [46, 47]. The rationale of approximating the transitions between the normal and cancer cell to simple switches are based on the following two facts.

1. The cancer process often exhibits switch-like behavior between two steady cell states, in accordance with the landscape-switching model. As a dysregulated cell developmental process [48], cancer progression is often determined by the regulation of the bistable gene expression states [49, 50], which correspond to the normal and cancer cell. Previous theoretical studies using a simple circuitry of the bistable switch between distinct gene expression states to study the cancer development well captured many characteristics of the cancer process [51, 30, 31]. The successes of these simplified models are likely due to the fact that the cell-fate decision-making processes can often be approximated to the transitions of the bistable switch [52, 53, 54].
2. From the physical perspective, cell cancerization and reversion are non-equilibrium non-adiabatic processes, which can be described by the landscape-switching model. The cancer progression requires extensive energy supplies that break the detailed balance of the system, leading to non-equilibrium dynamics. Furthermore, the transitions between the normal and cancer cell are impossible to occur spontaneously, so the waiting time of the inter-landscape hopping is expected to be much longer than the timescale of the intra-landscape dynamics. This feature of relatively slower inter-landscape dynamics compared to intra-landscape dynamics signifies a non-adiabatic process [55, 56]. In analogous to the surface hopping method [57], we separated the simulations of adiabatic chromosome dynamics at one cell state and non-equilibrium non-adiabatic inter-state switching dynamics, giving rise to the landscape-switching model.

We used the landscape-switching model to study the chromosome structural dynamics during the transitions between the human normal and cancer lung cell. We predicted that the chromosome can form more expanded structures than those at the normal and cancer cell during both of these processes but with different structural characteristics. Further analyses revealed that the chromosome at the transient intermediate state with a more expanded structure than those at the normal and cancer cell during the cancerization process share significant structural similarity to the one at the embryonic stem (ES) cell. This feature implies forming a cell state with the features of stemness or the CS cell during the cancerization process. We observed the high irreversibility for cancerization and reversion through the quantified structural transition pathways, the spatial repositioning of chromosomal loci and the hysteresis loop of establishing the chromosomal contacts. These findings underlined distinct mechanisms for these two cancer cellular processes. The prediction of forming a stem-like intermediate cell state during cancerization can be tested by the future time-course Hi-C experiments designed for the cancer developmental processes.

## 2 RESULTS

### 2.1 Chromosomal structural transition during cancerization and reversion

We used the landscape-switching model to investigate the chromosome dynamics during the cell cancerization and reversion processes. The implementation of the model is briefly summarized as follows. First, we iteratively fit a generic polymer model to reproduce the experimental Hi-C data through the maximum entropy principle simulation for the terminally differentiated human lung fibroblast cell (IMR90) and lung cancer cell (A549) (Figure S1), separately [59]. Previous studies have shown that the resulting potential not only captures the thermodynamics of the chromosome (i. e., Hi-C) [60, 61], but also describes the correct kinetic properties of the chromosomal loci diffusion within one cell state (one phase in cell cycle or one cell state in cell differentiation/reprogramming) [39, 40]. Here, we term the potential *V*(***r***|*S*) as the effective energy landscape for describing the chromosome dynamics in the normal or cancer cell, where ***r*** is the coordinate of the system at the cell state *S* (IMR90 or A549). Next, the simulation was set up with the chromosome exploring the structural dynamics under either the energy landscape of the normal (*V*(***r***|*IMR*90)) or cancer (*V*(***r***|*A*549)) cell. Then, the energy landscape underwent a switch from normal to cancer cell (*V*(***r***|*IMR*90) → *V*(***r***|*A*549)) or cancer to normal cell (*V*(***r***|*A*549) → *V*(***r***|*IMR*90)) to trigger the chromosomal structural transition during cancerization or reversion, respectively. Finally, the chromosome dynamics was governed by the post-switched energy landscape of the cancer cell (*V*(***r***|*A*549)) or normal cell (*V*(***r***|*IMR*90)). The model allowed us to observe the chromosome transformation during the cancerization and reversion processes with affordable computational resources (Figure 1).

**Figure 1:**
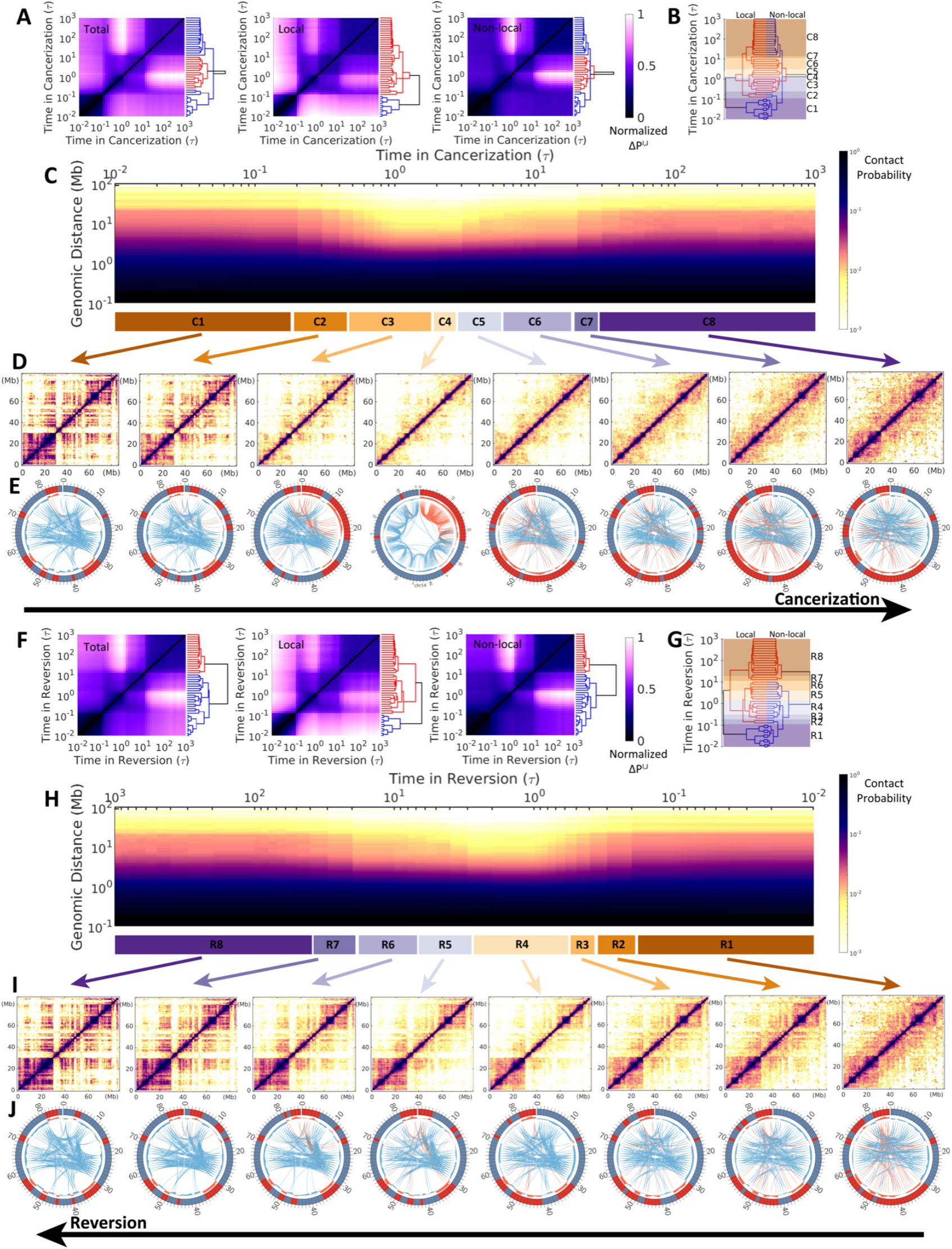
Chromosomal structural transitions during cancerization of IMR90 (normal cell) to A549 (cancer cell) *(Upper)* and reversion of A549 to IMR90 *(Lower).* (A) Hierarchical clustering of the chromosomal contact probability among each time frame *t* = I, *J* during cancerization varied by total, local (< 2Mb), and non-local (> 2Mb) contact ranges. The contact map difference is calculated with expression: 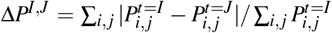, where 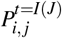 is the contact probability between chromosomal loci *i* and *j* at the time *t* = *I* or *J*. Δ*P^I,J^* is then normalized. (B) Reduced 8 stages (“C1”-”C8”) for the cancerization process based on the combination and comparison of the dendrograms of local and non-local Δ*P^I,J^* established in (A). Thus the chromosomes within one stage possess relatively similar contact probability maps. (C) The change of contact probability *P*(*s*) versus genomic distance *s* in the chromosome during cancerization with the 8 stages indicated at the bottom. (D) Hi-C heat (contact probability) maps of the chromosome for the 8 stages during cancerization. (E) The circle plots of chromosome for the 8 stages during cancerization [58]. The red and blue bands indicate the loci in compartment A and B, respectively. The compartment profiles are further shown as histograms near to the band plots. The connections show the long-range (> 5Mb) interactions which are identified by the *P_obs_/P_exp_*, where *P_obs_* and *P_exp_* are observed and expected contact probability, respectively [15]. The red, blue, and grey lines indicate the interactions between chromosomal loci from within compartment A, within compartment B, and between compartment A and B. The line widths correspond to the logarithmic scale of the *P_obs_/P_exp_* and only the top 200 weighted contacts are shown for better visualization. (F-J) are similar with (A-E) except for the reversion process of A549 to IMR90. Another reduced 8 stages during the reversion (“R1”-”R8”) are determined in (G).

We performed hundreds of independent landscape-switching simulations starting from different chromosome structures in the normal (and cancer) cell state to investigate the chromosomal structural transitions during the cancerization (and reversion) process (See “**Materials and Methods**”). By comparing the similarity of the contact probability formed through the pairwise chromosomal loci between time-series frames during the cancerization process (Δ*P^I,J^*, where *P* is contact probability and *I* and *J* are the time points), the transition can be reduced into two clusters according to the hierarchical clustering of Δ*P^I,J^* along with the processing time (Figure 1A). However, the clustering dendrograms for the local and non-local chromosomal contacts are very different, indicating distinct behaviors of structural rearrangements at the local and non-local ranges. The separation of the two clusters at the local contacts occurs early at ~ 0.1τ, where *τ* is the time unit in the simulation. In contrast, the non-local chromosomal contacts at the beginning of the transition are relatively similar to those at the late stages, and they are separated by the second cluster at ~ 0.5-10τ. This is also reflected by the contact probability *P*(*s*) versus the genomic distance s between a pair of chromosomal loci (Figure 1C), where an apparent decrease of P(s) is observed at time ~ 0.5-10τ.

By examining the clustering dendrogram patterns of the local and non-local contact formations in the chromosome, we further reduced the cancerization process into 8 stages (Figure 1B). As a result, chromosomes within one stage are structurally similar in terms of the local and non-local contacts. The contact maps of the 8 stages show a picture of how chromosomes perform the structural rearrangements in the caner formation process (Figure 1D). Interestingly, we found that as the can-cerization proceeds, the chromosome heat maps show lower probabilities (more sparse contacts in heat maps) than those in both normal and cancer cell. The result implies that the chromosome structure is expanding during cancerization. Besides, we observed a complex manner of the chromosome in organizing the compartment change during cancerization shown by the colored bands in the circle plots (Figure 1E). From the stage “C1” to “C3”, the populations of the chromosomal loci in compartment A increase associated with an increasing quantity of contacts (lines in each circle plot) formed within compartment A. At the stage “C4”, the chromosomal loci are roughly assigned to compartment A and B by sequence and the long-range contacts are rare. This implies the extensive breaking of the contacts formed in the chromosome at the normal cell. As cancerization further proceeds, a significant compartment-switching occurs during the stage “C4” to “C5”. From the stage “C5” to “C8”, the chromosomal compartment profiles and contact patterns gradually adapt to those at the cancer cell. Finally, we see that the chromosome in the cancer cell has more chromosomal loci in compartment A and forms more sparse contacts than the one in the normal cell. In the ES cells, chromosomes possess high populations of euchromatin, which maps with compartment A, to benefit the cell pluripotency and differentiation [62, 63, 64]. Our observation of the chromosome with populated expressive loci and open structure in the cancer cell resonates with the experimental evidence that the cancer cells possess certain contents of the “stemness” in favor of the self-renewal, differentiation, and proliferation [23, 65, 66].

Then we applied similar analyses into the reversion process. We found that the chromosomal structural transition during the reversion process is not the simple reversal of the cancerization. From the chromosomal contact formation perspective, the reversion process can also be grouped into two clusters (Figure 1F). However, these two clusters are separated by the time in sequence from the dendrogram plots. In detail, the time for separating the two clusters at local contact (~ 0.1τ) occurs much earlier than that at non-local contacts (~ 10τ). This indicates that chromosome structurally evolves faster at local than non-local range during reversion. Interestingly, the time evolution of P(s) ~ s also shows a chromosomal structural expansion at the time ~ 0.5-10τ (Figure 1H). However, the expansion appears to be less significant than that observed during cancerization. The chromosomal contact maps show distinct evolving ways deviated from the reversal of that in cancerization (Figure 1I). In particular, there is no significant compartment-switching during the reversion process, as the compartment undergoes gradual and moderate change, associated with reducing the contacts formed within compartment A (Figure 1J). Overall, we found that the chromosomal structural transitions undergo initial expansions followed by compactions during both the cancerization and the reversion processes but through irreversible paths.

### 2.2 Chromosome structural expansion during cancerization and reversion

To examine the chromosomal structural expansion during can-cerization and reversion, we monitored the contact probability *P*(*s*) versus the genomic distance *s*. *P*(*s*) provides information on how the chromosome organizes its structure along the genomic sequence, thus it dictates the polymer state of the chromosome [67, 68]. Chromosomes in both the normal and cancer cells show higher *P*(*s*) plots, corresponding to more compacted structures, than the one in the ES cell [64] (Figure 2A). This can also be inferred by the slope in the logarithmic relation of *P*(*s*) ~ *s* (Figure 2B). As the normal cell converts to the cancer cell, the profile of *P*(*s*) (colored by time) initially goes down and approaches the one at the ES cell (blue line), then increases to the cancer cell (purple line). The result draws a picture of the chromosome expanding its structure followed by compaction during cancerization. To quantitative measure this trend, we calculated the slope in the logarithmic relation of *P*(*s*) ~ *s*. We see that the slope at ~ 2τ has the lowest value and approximates the value found at the ES cell. This implies that the chromosome with the maximum structural expansion possesses a similar contact formation scaling versus the genomic sequence with the one at the ES cell. In other words, the chromosome may form the chromosomal structural characteristics of the ES cell during the cancerization. Besides, it is worth noting that the chromosome in the cancer cell is more expanded than it is in the normal cell, as the *P*(*s*) plot for the cancer cell is slightly below the normal cell.

**Figure 2:**
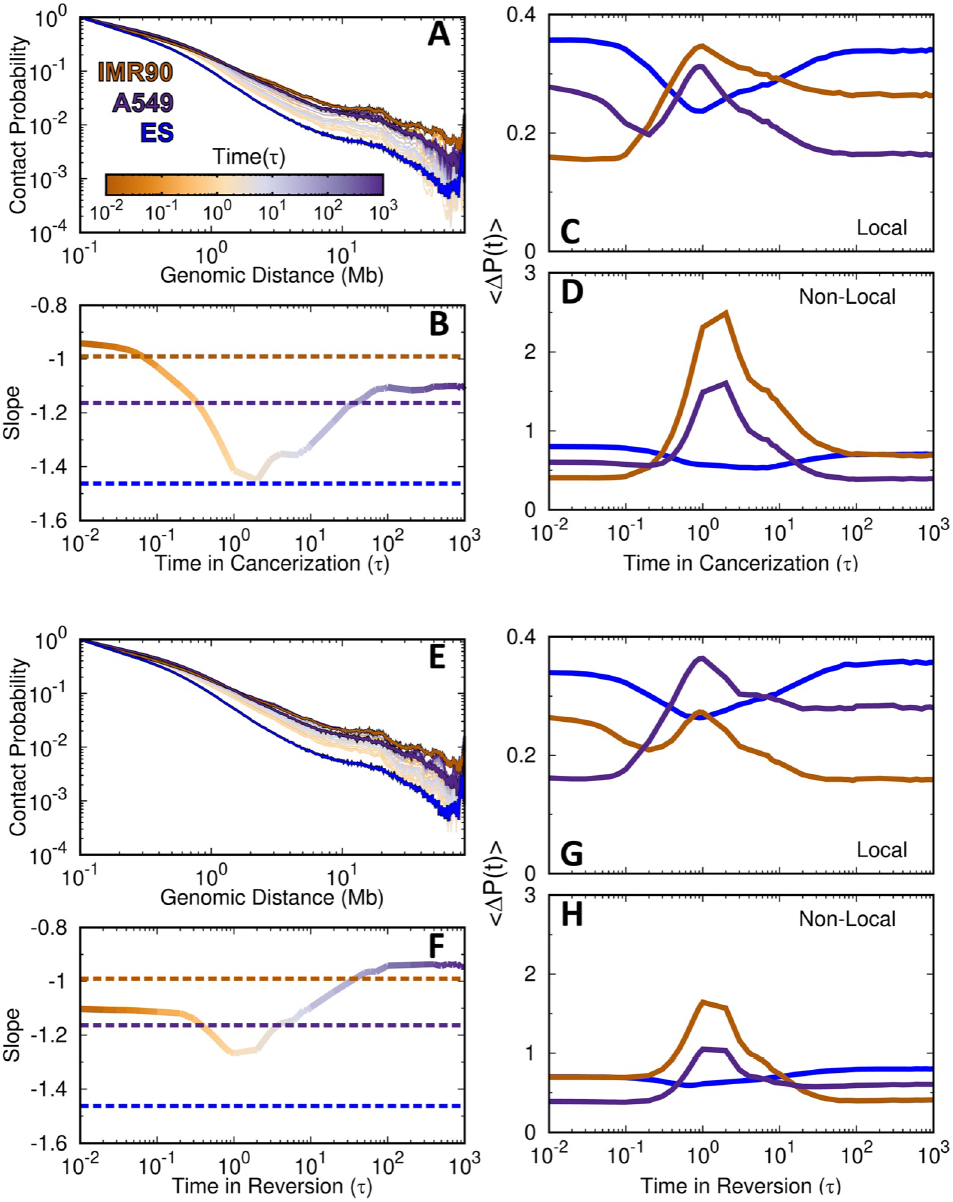
The time evolution of the chromosomal contact probability during cell cancerization and reversion. (A) The time evolution of contact probability *P*(*s*) versus genomic distance *s* during cancerization. The Hi-C data of IMR90, A549, and ES cells are also plotted. (B) The time evolution of the slope in the logarithmic relation of *P*(*s*) ~ *s* at the range of 0.5-7 Mb during cancerization. The differences in (C) local and (D) non-local 〈Δ*P*(*t*)〉 relative to the Hi-C data of IMR90, A549, and ES cells. (E-H) are the same with (A-D) except for the reversion process.

To measure the similarity of the chromosomal contact formation with that in the ES cell during cancerization, we calculated the average of the contact probability similarity 〈Δ*P*(*t*)〉 between the processing state at time point *t* during cancerization and the normal, cancer and ES cell at the local and nonlocal ranges (Figure 2C and 2D). We found that the chromosome at ~ 0.5-10τ, when the chromosome appears to be more expanded than those at the normal and cancer cell, as indicated in Figure 2B, 〈Δ*P*(*t*)〉 of the normal and cancer increase significantly while 〈Δ*P*(*t*)〉 of the ES cell decreases. This implies that the contact formation in the chromosome with more expanded structure than those at the normal and cancer cell, becomes similar to the one at the ES cell. The results indicate that the chromosome may go through stem-like structures to accomplish the cancerization process.

In contrast, the chromosome expansion in the reversion process from the cancer to the normal cell is not significant (Figure 2E). When the chromosome has the most significant structural expansion with the lowest *P*(*s*), the profile of *P*(*s*) still visu-ally deviates from the one in the ES cell. The steepest slope in *P*(*s*) ~ *s* shows a slight decrease from that in the cancer cell and occurs at the time ~ 1-3 τ. Besides, we can still observe that the similarity of the chromosomal contact formation between the processing state and the ES cell increases, but it is not as much as that observed during cancerization (Figure 2G and 2H). Therefore, the chromosome during the reversion process may possess less structural properties at the ES cell than during cancerization.

### 2.3 Quantified chromosomal structural transition pathways during cancerization and reversion

In order to obtain a quantitative picture of how the chromosomal structural transition occurs during the cancerization and the reversion processes, we projected all the transition trajectories as well as the averages onto several order parameters, which describe the shapes of the chromosome structure and contact formation at various ranges (Figure 3 and Figure S4). We first used the extension lengths of the chromosome structure along the longest and shortest principal axes (PA1 and PA3) [61] (Figure 3A). The individual pathways show high stochasticity, reminiscent of the stochastic dynamics in cell development [69]. During both the cancerization and reversion processes, there is an increase followed by the decrease of shape extension on the chromosome structure along with both PAs. However, from the average pathways, we see that the transitions of these two processes do not follow the same routes. This is also observed by projecting the pathways onto the radius of gyration (*R_g_*) and aspheric quantity (Δ). Δ measures the asphericity of the chromosome structure, and the deviation of Δ from 0 measures the deviation from the perfect sphere [70]. Besides, we can see that the shape of the chromosome with the most significant expanded structure is geometrically different from the one in the ES cell. However, the chromosome at the stage “C4” in cancerization is closer to the ES cell than the stage “R4” in reversion.

**Figure 3:**
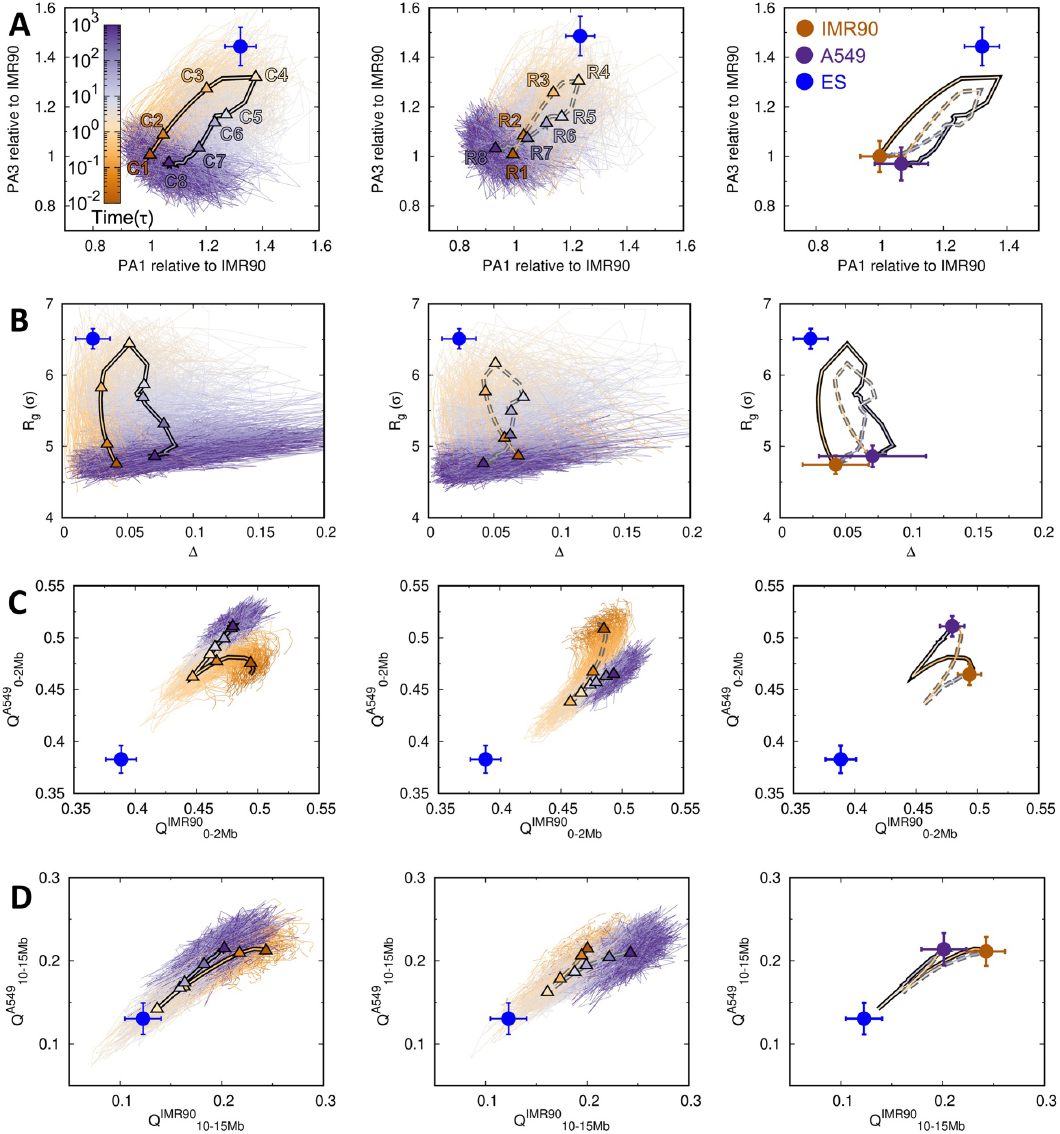
The pathways of chromosomal structural transitions during cell cancerization and reversion. The pathways are projected onto the order parameters of the chromosome with all trajectories presented for cancerization *(Left)* and reversion *(Middle).* The 8 stages are indicated by triangles (“C1”-”C8” in cancerization, *Left)* and diamonds (“R1”-”R8” in reversion, *Middle)* on the averaged pathways of the cancerization (solid line) and reversion (dashed line), respectively. The averaged pathways are additionally shown and colored by time *(Right).* The quantities of the chromosome in the IMR90, A549, and ES cells are correspondingly placed as brown, purple and blue points, respectively. The pathways are projected onto (A) the extensions of the longest and the shortest principal axes (PA1 and PA3), (B) the radius of gyration *(R_g_*) and the aspheric quantity (Δ), contact similarity in terms of the fraction of native contact *Q* to the IMR90 and A549 at (C) local range (0-2Mb), and (D) long-range (10-15Mb). The data of the ES cell are obtained from our previous work [45].

The chromosome structural expansion during the cancerization and reversion can also be reflected by the local and nonlocal contact formations (Figure 3C and 3D). Here we used the fraction of “native” contact *Q*, which was widely applied in protein folding [71], with references being the average pairwise distances between loci in chromosome ensembles at the normal and cancer cells. There are different pathways for local chromosome structural formations during these two processes, which appear to deviate significantly from the chromosome in the ES cell. For non-local contacts, the pathways become overlapped but still show differences at the most expanded chromosome structure. The results imply that the highly irreversible chromosomal structural transition occurs universally at both the local and non-local range.

### 2.4 Spatial rearrangement of chromosomal loci during cancerization and reversion

It has been recognized that the spatial distribution of the chromosomal loci strongly influences the transcriptional activity [72, 73]. To see how the chromosome rearranges the spatial distribution of the chromosomal loci along with the compartment formation during cancerization and reversion, we calculated the radial density *ρ* (*r*) of chromosomal loci and further classified it into the compartment A and B. We found that the normal cell, which is a terminally differentiated cell, tends to locate the chromosomal loci of compartment A and B towards the chromosome’s surface and interior, respectively (Figure 4A). The finding is in line with previous simulations [74, 40, 75] and experiment [76]. In contrast, the chromosome in the ES cell exhibits a roughly uniform distribution of the chromosomal loci regardless of the compartment states. For the cancer cell, *ρ* (*r*) appears to be the intermediate between those at the normal and ES cell. We further calculated the radial density similarity Δ*ρ* (*t*) between the processing state at time point *t* and the normal, cancer and ES cell for the total loci and the loci in compartment A and B (Figure 4B). When the normal cell converts to the cancer cell, Δ*ρ* (*t*) of the ES cell decreases and reaches the minimum at time ~ 0.9-3τ, which are varied by the total loci and the loci in the compartment A and B (Figure 4B). We note that this period corresponds to the stage when chromosome forms more expanded structure than those at the normal and cancer cell (Figure 2B), so the results indicate that the spatial distribution of the loci in chromosome during this period is similar to that in the ES cell. Further proceeding cancerization decreases Δ*ρ* (*t*) of the cancer cell. Overall, we found a non-monotonic spatial repositioning of the chromosome loci during cancerization. In the beginning, the chromosome segregates the active loci located in compartment A and inactive loci located in compartment B towards the surface and interior, respectively (Figure 4C). Then, the compartment segregation of the chromosome is still effective, but it forms a stem-like pattern with a uniform radial distribution of genomic loci regardless of the compartment states. Finally, the spatial distribution of the genomic loci reaches the one at the cancer cell.

**Figure 4:**
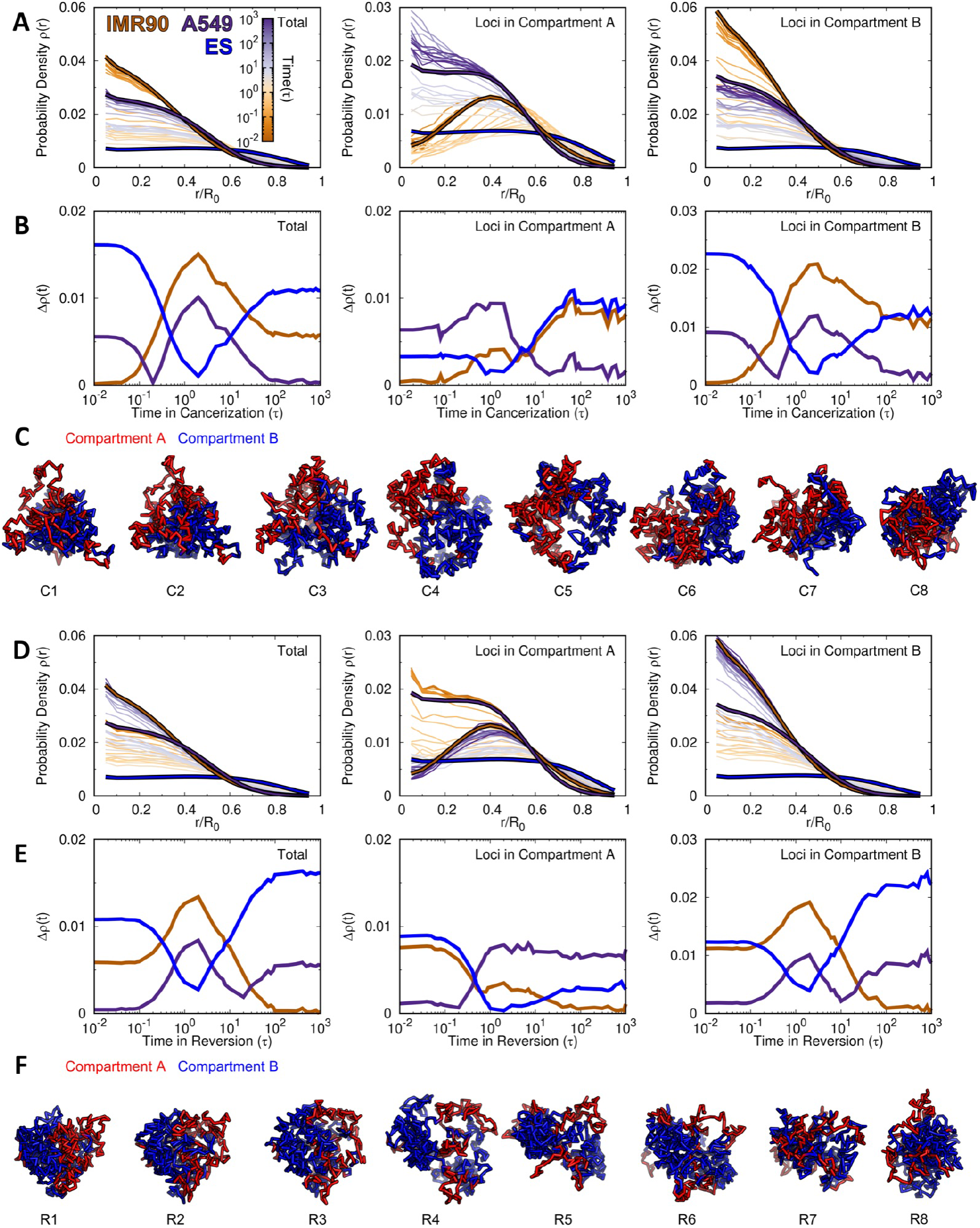
The change of the radial density in the chromosome during cell cancerization and reversion. (A) The change of the radial density profile *ρ* (*r*) for the whole loci *(Left),* loci in compartment A *(Middle)* and loci in compartment B *(Right)* in the chromosome during cancerization. The profiles of the IMR90, A549 and ES cells are colored brown, purple and blue, respectively. (B) The difference of the radial density, calculated by 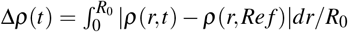, from the processing time (*t*) to the reference cells (Ref). (C) Structural illustrations of the chromosome with loci colored by compartment states during cancerization. (D-F) are the same with (A-C) except for the reversion process.

Interestingly, we observed a similar trend of spatial repositioning of the chromosomal loci during the reversion process (Figure 4(D-F)). In other words, *ρ* (*r*) during reversion also goes through processing states sharing similarity with that in the ES cell. In detail, we found that Δ*ρ* (*t*) of the ES cell decreases more for the loci in compartment A than the ones in compartment B at time ~ 1-3τ (Figure 4E), corresponding to the stage when chromosome forms more expanded structures than those at the normal and cancer cell (Figure 2F). This is different from that observed in the cancerization process, where spatial repositioning of the chromosomal loci in compartment B is dominant to be the stem-like, as decreasing Δ*ρ* (*t*) of the ES is more significant for the loci in compartment B than the ones in compartment A. The combined results suggest that during both cancerization and reversion processes, the chromosome dynamically rearranges the spatial positions of the chromosomal loci and the distribution can resemble the one at the ES cell.

### 2.5 Stemness and irreversibility of chromosomal structural transitions during cancerization and reversion

Our results have implied that the chromosome during the can-cerization and reversion may adopt the chromosome structure in the ES cell. To quantitatively assess this, we divided the contacts by different ranges and then performed the PCA on the contact probability evolving trajectories. Since the contacts at long-ranges are usually formed with very low probabilities, it may introduce large uncertainty and imprecision when directly applying to PCA. To resolve this issue, we instead used the matrix *P_obs_/P_exp_*, which captures the essence of the compartment formation [15]. We also note that the boundaries of TADs are mostly conserved during the transitions (Figure S5). This indicates that the chromosomal structural arrangement during can-cerization and reversion should mostly rely on the contacts at the long ranges, which can be captured by the compartment.

We projected the transitions onto the first and second most weighted PCs (Figure 5). In this respect, at the local range (< 2 Mb), we see that the ES cell is not on either of the two pathways. As the contact range increases (2-5 and 5-10 Mb), the location of the ES cell becomes close to the cancerization pathway but is far from the reversion pathway. Further increasing the contact range to 10-15 and 15-20 Mb shows that the ES cell is close to the cancer cell. When the contact range is very long (20-40 Mb and > 40 Mb), the ES cell locates far from both of these two pathways. Our results showed that the chromosome during the cancerization process may explore the structures formed in the ES cell at the moderate contact range (2-20 Mb). In contrast, the chromosome during the reversion process seems to be structurally different from the one in the ES cell, though the chromosome is prone to adopt many of the structural properties in the ES cell, such as the contact probability scaling, structural shape geometry and spatial distribution of the chromosome loci.

**Figure 5:**
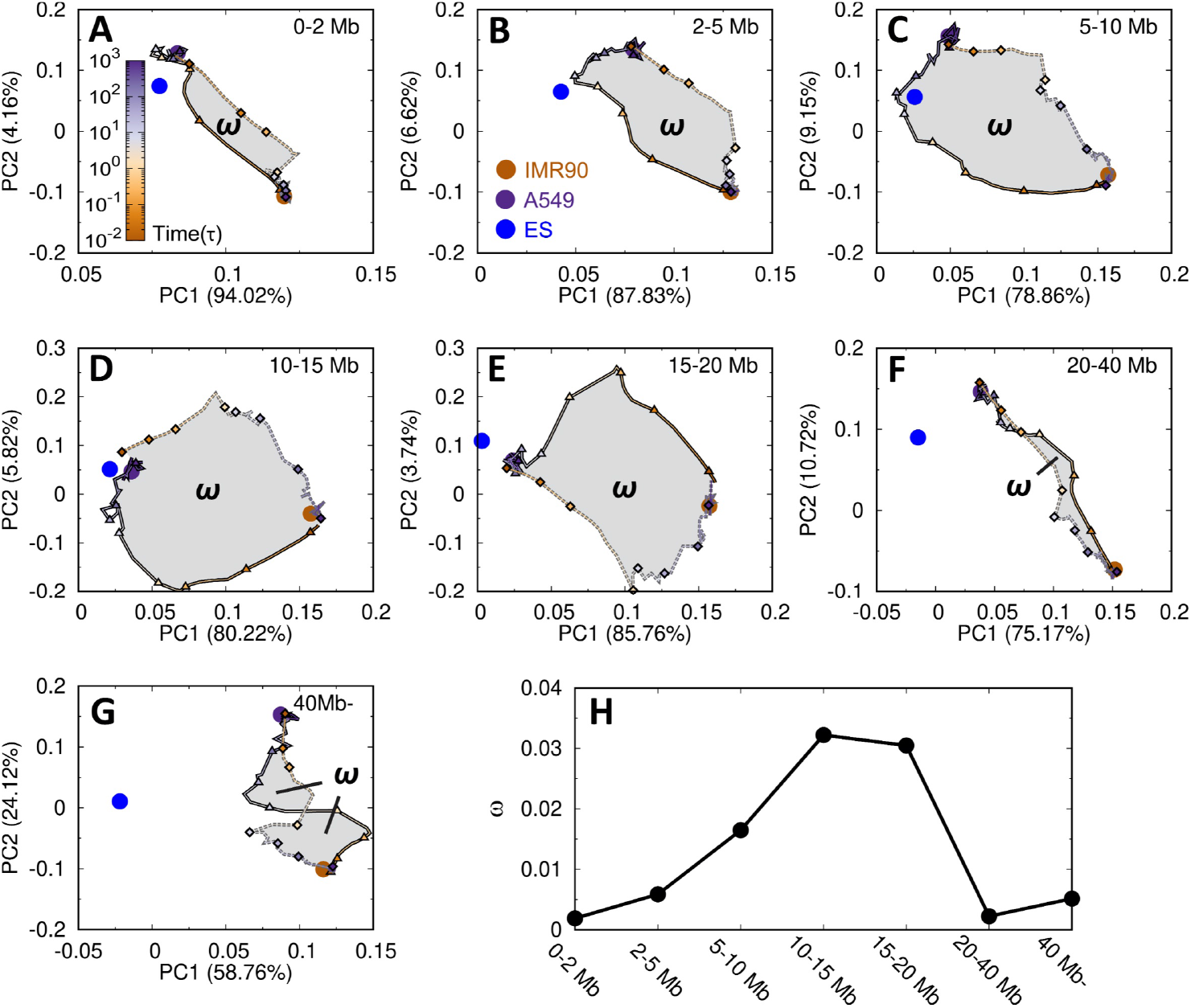
The pathways of the chromosomal structural transition during cell cancerization and reversion. The principal component analysis (PCA) of the contact matrix *P_obs_/P_exp_*, which is further divided into different ranges (A-G). The reference points from the IMR90, A549 and ES cells are plotted as brown, purple and blue points, respectively. The hysteresis loop (ω) is colored grey in (A-G), and the area is calculated for different contact ranges (H).

Interestingly, we found that the two pathways do not overlap, resulting in the hysteresis phenomenon. Hysteresis indicates a toggle-like switching behavior and was often described in the context of ferromagnetism. In biology, hysteresis was found in the cell cycle and demonstrated as the driving force for the irreversible cell-cycle transitions [77, 78]. In this respect, the hysteresis loop area provides a quantitative measure of the degree of the irreversibility, thus it was calculated here (Figure 5H). We found that the largest hysteresis area, which corresponds to the most significant irreversibility of the chromosomal structural transitions during cancerization and reversion, is at the moderate contact range. Combining the results, we showed that the chromosomal structural transitions during the cancerization and reversion processes are highly irreversible, and the chromosome tends to form stem-like structures during the cancerization.

## 3 DISCUSSION AND CONCLUSIONS

We used the landscape-switching model to explore the chromosomal structural transition during the cell cancerization and reversion processes. More than two hundred independent simulations (226 simulations for cancerization and 246 simulations for reversion, see **Materials and Methods**) were performed for a long time under the potential designated for describing the chromosome dynamics in the normal (or cancer) cell state. We found that these simulations with the initial chromosome structures chosen based on the clustering method recapitulate the structural ensembles in terms of the contact maps, TADs and compartments (Figure S6). The feature suggests that the selected chromosome structures can represent the chromosome structural distributions at the original cell states and are statistically sufficient to initialize the landscape-switching simulations.

Our structural analyses showed that the chromosome with the highest degree of structural expansion during cell cancerization and reversion share significant structural similarity to the chromosome at the ES cell in many aspects, including the trend of contact probability declining with the genomic distance, the global structural geometry and the spatial distribution of chromosome loci. However, there are differences between the chromosomes with expanded structures during these two processes. As shown by the enhanced contact probability evolving trajectories (Figure 5), the chromosome during can-cerization can form a stem-like chromosome contact pattern up to the range of 20 Mb. In contrast, this is not observed during reversion. The results suggested a potential role of the cell state with stemness during cancerization in guiding the process.

Based on our simulation results, we can propose a pictorial Waddingtons landscape from the chromosomal structural perspective to understand the cancerization and reversion (Figure 6). In the context of Waddingtons epigenetic landscape [79], the cell, metaphorically referred to as a ball, rolls down from the ES cell at the top, which possesses the highest degree of stemness, to the terminally differentiated cell at the basin of the land scape to accomplish cell differentiation. As the reverse process, cell reprogramming transforms the differentiated cell to the ES cell by gaining the stemness [80, 81]. Hi-C data revealed that the chromosome at the cancer cell has more loci in compartment A than compartment B (Figure 1), corresponding to more populated active, open euchromatin than that of inactive, closed heterochromatin. This feature suggests that the cancer cell, which exhibits the characteristics of the stem cell, should be located at a landscape layer, higher than the normal cell, which is terminally differentiated with a large proportion of the chromosomal loci in repressive compartment B (Figure 1). Transforming the normal cell to cancer cell increases the stemness. However, our simulations showed that cancerization is a non-monotonic process, during which the chromosome with the highest degree of structural expansion exhibits significant structural similarity with the one at the ES cell. In the landscape view, the cell at the first stage of can-cerization climbs up on the landscape approaching the ES cell, reminiscent of cell reprogramming. The second stage of cell cancerization corresponds to a rolling down process toward the cancer cell, reminiscent of cell differentiation. On the other hand, the chromosome during the reversion process also expands and reorganizes its structure towards the one at the ES as an implication of gaining the stemness. However, the structural difference of the chromosome to the one at the ES during reversion is more significant than that during the cancerization. This leads to a relatively weak reprogramming process at the first stage of the reversion process. Thus the reversion pathway should appear to be under the cancerization pathway (Figure 6).

**Figure 6:**
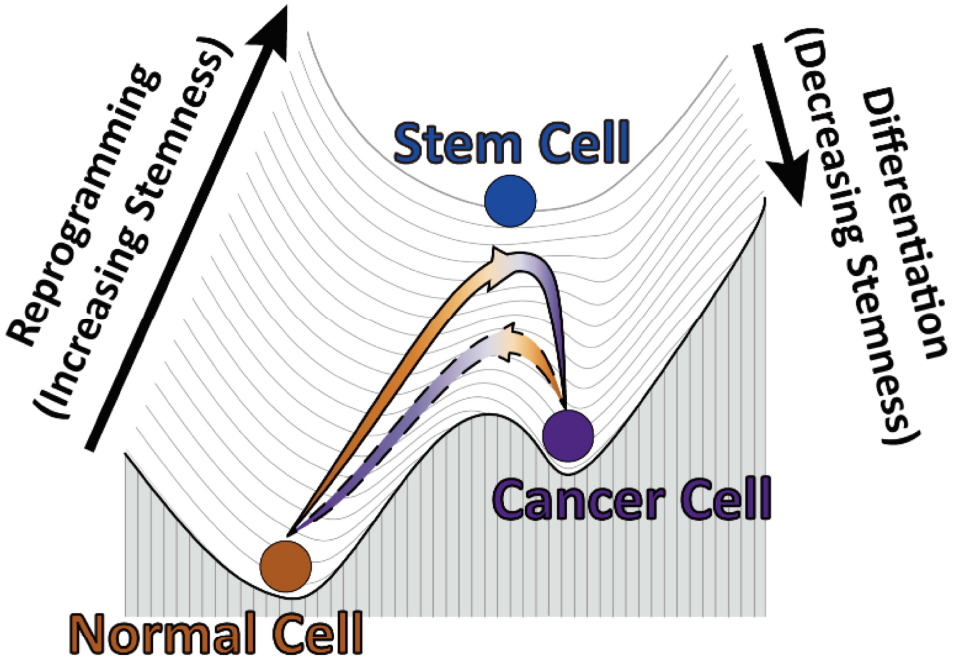
The Waddington landscape of the cell cancerization (solid line) and reversion (dashed line) processes from the chromosomal structural transition perspective.

Our simulation showed that cancerization and reversion are irreversible processes. Previously, it was assumed that the forward and reverse processes of cell state transitions may pass through the same intermediate cell states, referring to as the metastable attractors on the Waddingtons landscape [82, 83]. Theoretical studies at the gene network level have shown that irreversibility is a universal feature for cell state transition due to the non-equilibrium effects [84, 85]. Our simulation results provided a molecular-level description that the irreversibility of cell cancerization and reversion can be reflected by the chromosome structural dynamics. Besides, we proposed a quantitative way to measure the irreversibility in cancerization and reversion using the hysteresis loop analysis of the chromosome contact formation. Our findings also resonate with a recent experiment, which used a combination of Hi-C and replication timing analyses to characterize the irreversible non-overlapped pathways between the cell state transition, namely the differentiation and reprogramming [86].

The stemness-related genetic markers and interactions in cells are controlled by the underlying gene regulatory network [87], which uses chromosome as the structural scaffold for functionalities. Besides, the CS cells were found to have similar marker expression profiles with normal stem cells [88]. These features support that our approach for inferring the stemness and connecting it to the partial formation of the CS cell is reasonable and based on the intimate structure-function relationship at the genomic level [72, 4, 5]. Therefore, we speculate that the stem-like intermediate cell state during cell can-cerization may contribute to the formation of the CS cells, of which the terminally differentiated somatic cell has been suggested as one potential origin [89]. Due to the lack of the Hi-C data at the CS cells, the precise and direct comparisons of the intermediate state to the CS cells were not performed. A strict assessment of our statement can be made when the experimental data are available.

Our predictions of the transient intermediate state with stemness during cell cancerization may have implications for cancer treatments. The increase of the stemness during cancerization may promote the cell self-renewal and differentiation processes, similar to what the CS cells do. Thus, decreasing the stemness of the cell by deviating the chromosome structures from the one at the ES cell can help to repress the cancer initiation and progression. Further efforts can be focused on developing effective genome structure engineering methods for modulating and disrupting the specific chromosomal interactions similar to those formed in the ES cells [90,91,92,93,94], serving as a potential therapeutic approach.

Our predictions can be tested by future experiments. In this regard, the time-course Hi-C experiments appear to be promising and competent in monitoring the chromosomal structural evolution during the cancerization and reversion processes [95]. It is expected that the Hi-C data at the high temporal resolution can provide a dynamic picture of how the chromosome structure rearranges and offer useful clues as to the formation of the stem-like intermediate state during cell state transition using advanced data-analysis methods [96, 97]. Meanwhile, mounting theoretical and experimental studies confirm the existence of intermediates during the epithelial-mesenchymal transition (EMT) [98, 99, 100, 101], which contributes to cancer metastasis and tumor relapse. The EMT intermediates often exhibit a gain of stemness compared with the epithelial and mesenchymal cells [102, 103, 104, 105], leading to a non-monotonic increasing-followed-by-decreasing trend in stemness during the EMT [106]. These properties of the EMT process are similar to what we have observed in our simulations of the transition from the normal cell to the cancer cell. Therefore, the fruitful approaches developed for uncovering the EMT intermediate and studying the EMT pathways can be adapted to investigate the cancerization processes [107]. Ultimately, the combined efforts on the direct cancerization and the EMT processes will help us to complete the understanding of how cancer initializes and metastasizes.

In summary, we presented a chromosomal structural-level study of illuminating the molecular mechanisms of cancer formation and its reversion processes. The targeted cells belong to the human lung organism and should be regarded as one example of cancer. Up to now, more than a hundred distinct types of cancer have been uncovered. Different types of cancer may vary substantially in their behaviors. However, all types of cancer are caused by the common mechanism of uncontrolled cell division, reflecting the dysregulation of cell self-renewal. Therefore, we speculate that our observation of the stem-like intermediate state during cancer formation may likely be observed by the other types of cancer. Recent experimental observations that the chromosome opening its structure in the early stages of tumor formation is regardless of cancer types and shares a similar trait of chromosome organization with the stem cell provide evidence of our theoretical findings and speculations our speculation [108, 109]. The open chromosome structure in the stem-like intermediate state may contribute to the increased genomic instability and active transcription for cells to gain plasticity that has been demonstrated as the universal key to facilitate cancer progression [110]. Future studies on elucidating the cancer formation pathways and the stem-like intermediate state can help to decipher the fundamental mechanisms that underlie the different types of cancer.

## Materials and Methods

### Chromosome polymer model

The basic background of our polymer model is a generic polymer model with homogeneously weighted bonded and nonbonded potentials:

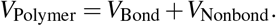

The bonded potential contains the finitely extensible nonlinear elastic (FENE) bond stretching interactions, linear-promoting angle bending interactions, and neighboring beads non-overlap interactions with the expression as follows [111]:

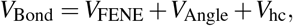

where the FENE potential for the bond (*r*_*i,i*+1_) between neighboring beads (i, i+1) is expressed as:

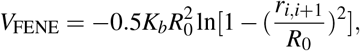

and the angle potential is applied to the angle (θ_i_) by the three adjacent beads with the following form:

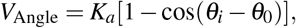

and the non-overlap interaction between (i, i+1) is described by a hard-core Lennard-Jones potential (*V_LJ_*):

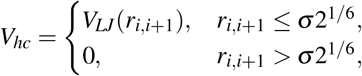

where

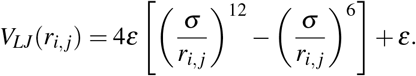

The nonbonded potential for every bead pair (i,j) uses the soft-core interactions that allow the chain-crossing to mimic the effects of topoisomerase [112].

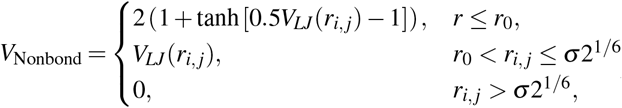

In addition, a spherical confinement was used to mimic the volume fraction of the chromosome in the cell nucleus at 10%, same as used previously [111, 60]. With only the generic polymer potential, the model generates an ensemble that resembles the equilibrium globule [45].

Reduced units were used throughout simulations. The energy unit was *ε* = 1.0. The bond length *σ* was set to be the length unit. In FENE potential, *R*_0_ = 1.5*σ* allows the bonds to stretch flexibly and *K_b_* = 30.0/*σ*^2^. The angle potential has a strength of *K_a_* = 2.0 and angle of *θ*_0_ = *π*. In the soft-core potential, we set 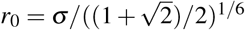, so *V_LJ_*(*r*_0_) = 2.0 [60]. The temperature in all simulations was set to 1.0 in energy units by multiplying by the Boltzmann constant. Langevin stochastic dynamics was applied with a time step of 0.001τ and a friction coefficient of 10.0τ^−1^, where τ is the reduced time unit.

### Maximum entropy principle simulation

The potential *V*(***r***) in the maximum entropy principle simulations is made up of the generic polymer potential *V*_Polymer_ and the Hi-C restraint potential *V*_Hi-C_:

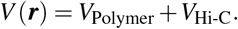

The entropy of system under *V*(***r***) relative to a given prior distribution *ρ*_0_(***r***) under polymer potential *V_Polymer_* is defined as [59]:

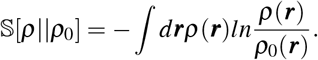

This entropy should be maximized subject to the following constraints in order to be compatible with observations (contact probability ***P**_i,j_* and experimental Hi-C data *f_i, j_*):

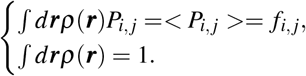

The maximization of the entropy can be obtained using the method of Lagrangian multipliers by searching for the stationary points of the Lagrange function:

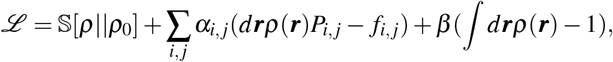

where *α_i,j_* and *β* are Lagrangian multipliers. By setting 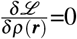 and neglecting the normalization factor, the posterior distribution has the following expression:

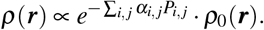

Therefore, the potential *V*(***r***) with maximum entropy principle, is expressed as follows:

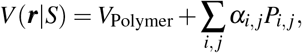

where *V*_Hi-C_ is in a linear form of contact probabilities and *S* represents the cell state (IMR90 or A549). In practice, *P_i, j_* is the calculated contact probability between chromosomal *i* and *j* using step function and *α_i, j_* is the corresponding contact strength, which is iteratively adapted by simulations. During the iteration process, the contact probability *P_i, j_* is restrained by the Hi-C data *f_i, j_*. In the end, the chromosome under the potential *V*(***r***|*S*) generates an ensemble that can reproduce the Hi-C maps at the normal and cancer cell, separately (Figure S1). Details of the maximum entropy principle simulation can be found in previous studies [60, 61]. The maximum entropy principles simulations for the ES and IMR90 cells were done in our previous work [45].

### Landscape-switching model

We used the resulting potentials *V*(***r***|*S*) of the maximum entropy principle simulations for the normal and cancer cells to represent the effective energy landscapes. We performed the hierarchical clustering on the chromosome ensembles generated by the maximum entropy principle simulations. Two chromosome structures in each cluster, which has a population higher than 0.3%, were picked out as the initial structures for performing the landscape-switching simulations (Figure S2). The purpose of choosing the chromosome structures in the populated clusters rather than using the whole sets of the structures in the ensembles is to generate a limited number of structures that can sufficiently represent the ensemble in order to significantly reduce the computational expenses. This led to 226 and 246 trajectories for the cancerization and reversion processes, respectively.

The landscape-switching simulations were performed as follows. First, the simulations were run under the potential *V*(***r***|*IMR*90) (cancerization) or *V*(***r***|*A*549) (reversion) for 5000τ, where τ is the time unit of the simulations. Then a sudden switch of potential in terms of *V*(***r***|*IMR*90) → *V*(***r***|*A*549) (cancerization) or *V*(***r***|*A*549) → *V*(***r***|*IMR*90) (reversion) was implemented. Finally, the simulations were under the new potential *V*(***r***|*A*549) (cancerizatin) or *V*(***r***|*IMR*90) (reversion) for 10000τ. All the trajectories were collected and combined for the analyses to generate the results present in this study.

## Acknowledgement

We acknowledge the support from the National Science Foundation PHY-76066. The authors would like to thank Stony Brook Research Computing and Cyberinfrastructure, and the Institute for Advanced Computational Science at Stony Brook University for access to the high-performance SeaWulf computing system, which was made possible by a $1.4M National Science Foundation grant (#1531492).

## Notes

### Competing Interest Statement

The authors have declared no competing interest.

