## Supporting Information for "Deciphering the molecular mechanism of the cancer formation by chromosome structural dynamics"

for

### 1 Materials and Methods

#### Hi-C data processing

We used the human fetal lung fibroblast cell (IMR90) and human lung carcinoma cell (A549) as the normal and cancer cells to investigate the cancerization and the reversion processes. The Hi-C data of the embryonic stem (ES) cell and IMR90 were downloaded from the publicly available Gene Expression Omnibus (GEO) repository archives with accession number GSE35156 [1] and the Hi-C data of A549 cell line was obtained from the ENCODE project [2] with GEO accession number GSE105600. All the replicas were combined and proceeded to the Hi-C Pro following standard pipelines for generating the contact maps at a resolution of 100kb [3]. In this study, we focused on the long arm of chromosome 14 (20.5-106.1Mb), so the polymer model has 857 beads.

#### Identifications of compartments

The compartment profiles were calculated by the enhanced contact probability $P_{obs}/P_{exp}$, which is the ratio between the observed contact probability $P_{obs}$ and expected contact probability $P_{exp}$ [4]. The enhanced contact probability map was built at a resolution of 1Mb. Then the Iterative Correction and Eigenvector Decomposition (ICE) method was used to perform the normalization [5]. The principal component analysis (PCA) was performed on the normalized matrix and the first principal component (PC1) was referred to as the compartment profiles. The direction of the PC1 values is arbitrary, and we set the positive and negative PC1 values with gene density (positive to gene-rich and negative to gene-poor). This was done on the IMR90 data. Then, the direction of the PC1 values during cancerization and reversion was determined according to the correlation coefficient with the PC1 values of the IMR90 data.

[4] Erez Lieberman-Aiden, Nynke L Van Berkum, Louise Williams, Maxim Imakaev, Tobias Ragoczy, Agnes Telling, Ido Amit, Bryan R Lajoie, Peter J Sabo, Michael O Dorschner, et al.

Comprehensive mapping of long-range interactions reveals folding principles of the human genome. *Science*, 326(5950):289–293, 2009.

[5] Maxim Imakaev, Geoffrey Fudenberg, Rachel Patton McCord, Natalia Naumova, Anton Goloborodko, Bryan R Lajoie, Job Dekker, and Leonid A Mirny. Iterative correction of hi-c data reveals hallmarks of chromosome organization. *Nature methods*, 9(10):999, 2012.

[6] Ruxandra I Dima and D Thirumalai. Asymmetry in the shapes of folded and denatured states of proteins. *The Journal of Physical Chemistry B*, 108(21):6564–6570, 2004.

### 2 Figures

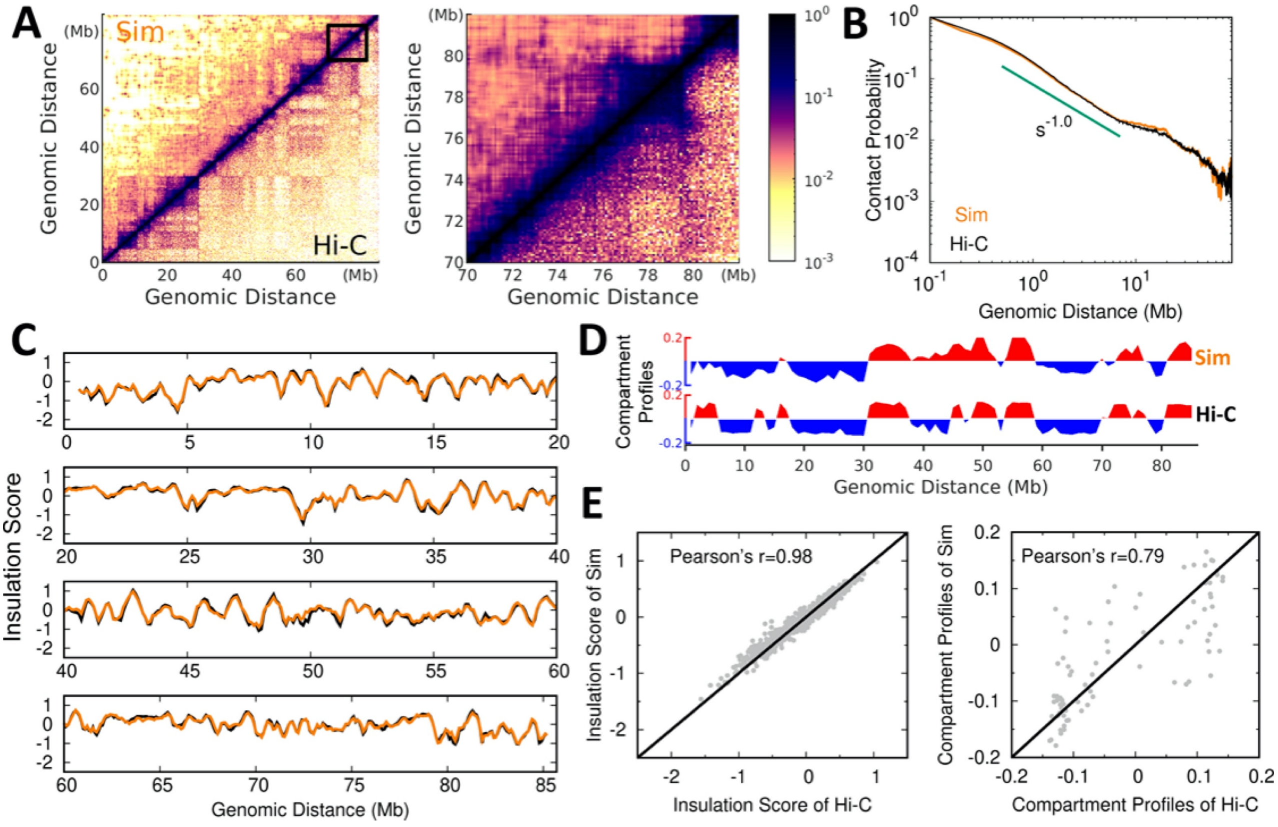


Figure S1: Comparisons of simulations and Hi-C data in the A549. (A) Hi-C contact maps of chromosome from maximum entropy principle simulations and experiments at global (*Left*) and local (*Right*) scales. (B) Contact probability versus genomic distance in chromosome for simulations and Hi-C data with a slope of -1.0 observed. (C) Insulation score of chromosome obtained by simulations and Hi-C data. (D) Compartment profiles of chromosome obtained by simulations and Hi-C data. (E) Correlations of insulation score (*Left*) and compartment profiles (*Right*) between simulations and Hi-C data.

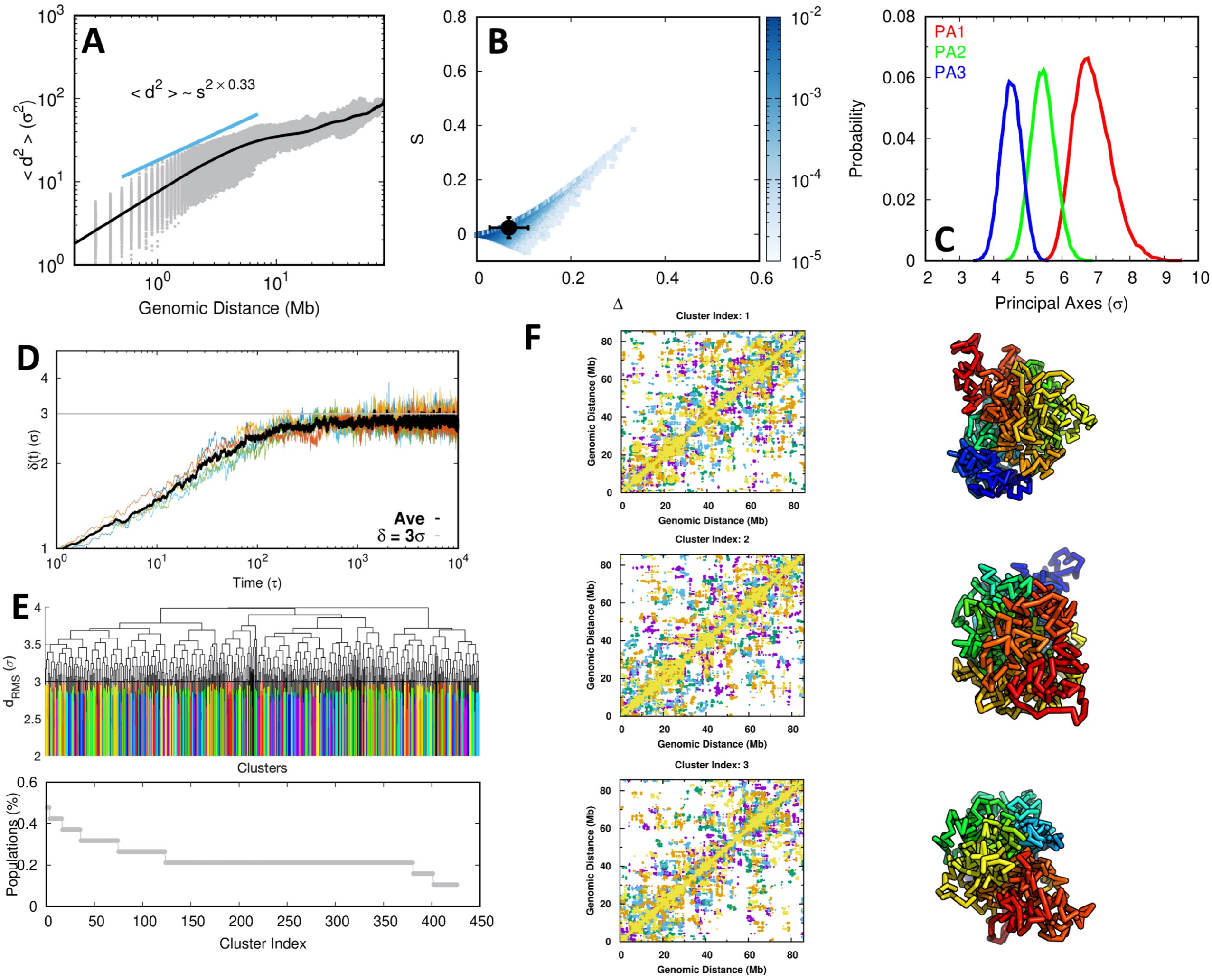


Figure S2: Chromosome ensemble in the A549. (A) Contact distance ($d$) versus genomic distance ($s$) in the chromosome. (B) The probability distributions of aspheric parameters of chromosome. $\Delta$ and $s$ are calculated using the inertia tensor [6]. Deviation of $\Delta$ from 0 (the value corresponding to a sphere) gives an indication of the extent of anisotropy. Negative values of $S$ correspond to oblate shapes and positive values to prolate shapes. (C) The probability distribution of configurational extension on three principal axes of the chromosome. (D) The time evolution of average root mean square distance ($d_{rms}$) between every genomic pair in chromosome at the time $t$ relative to its initial value: $\delta(t) = \sum_{i,j} d_{rms}(i, j, t)/N_{pairs}$, where $N_{pairs}$ is the number of summed pairs. The maximum of $\delta(t)$ is close to $3\sigma$. (E) The hierarchical clustering of the chromosome shown as a dendrogram (*Top*) and the populations of the cluster (*Bottom*). Cut-off distance $3\sigma$ was applied. (F) The top 3 most populated chromosome clusters. Each is shown with a mixed contact map (*Left*), which contains 5 structures within the cluster, and one representative structure (*Right*).

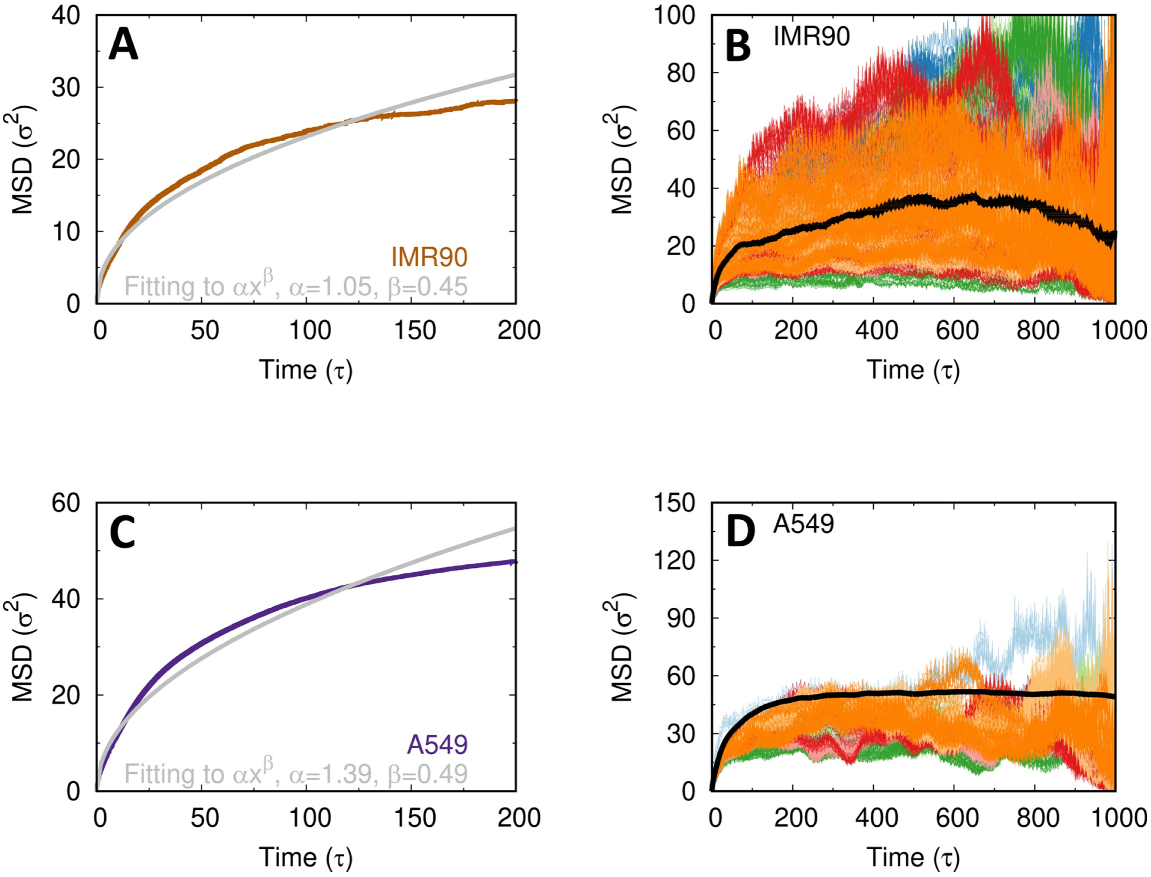


Figure S3: Chromosome diffusion dynamics in the IMR90 and A549. (A) Fitting of MSD to the power-law function (MSD $\sim \alpha x^{\beta}$) in IMR90. MSD is calculated by averaging the MSD from 5 independent simulations with the potential from the maximum entropy principle simulation. (B) MSD of all the individual chromosomal loci in the IMR90 obtained from one simulation with average shown as the black line. (C) and (D) are same as (A) and (B) but for the A549 cell.

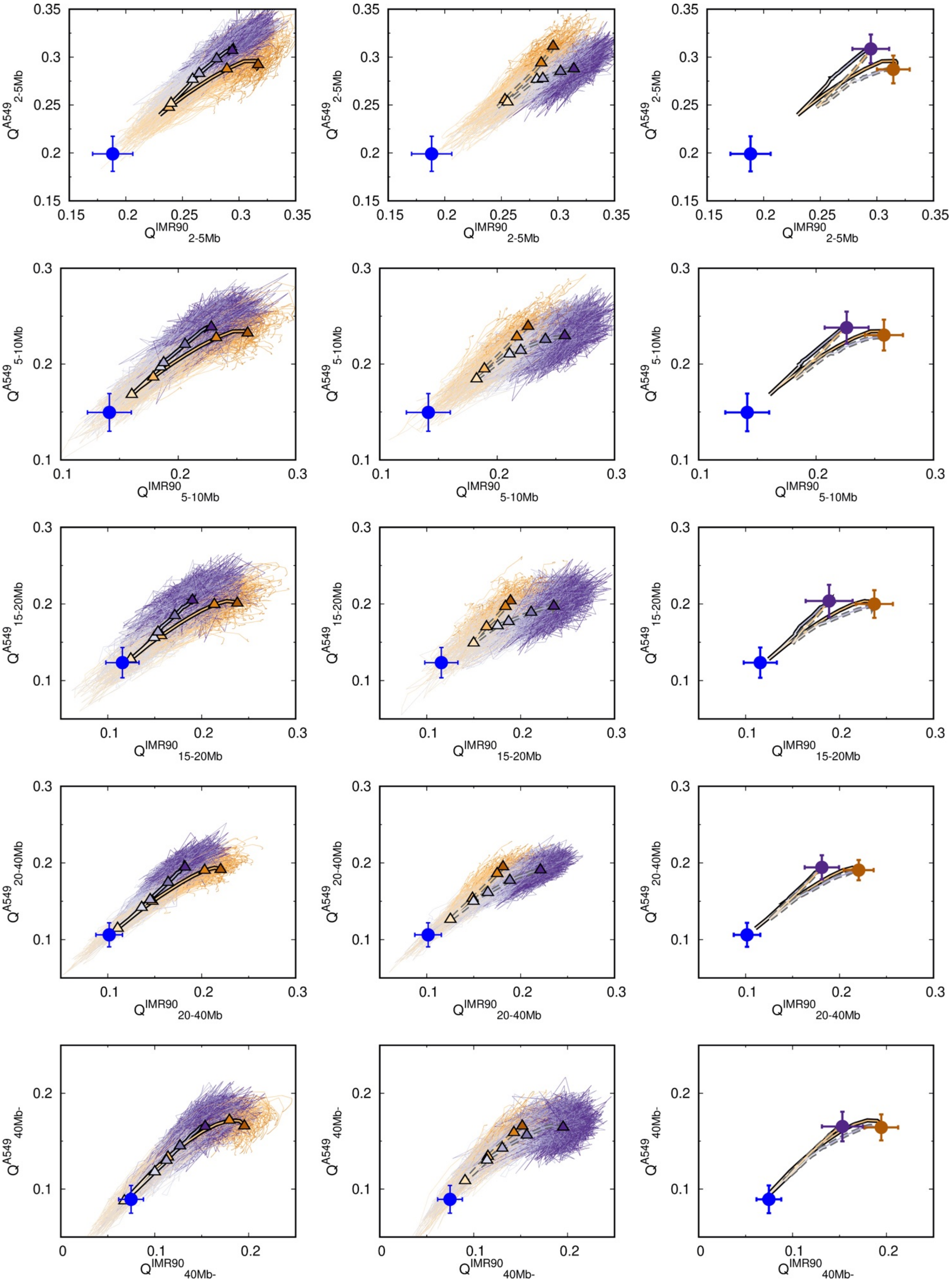


Figure S4: Pathways of chromosome structural transitions projected on $Q$ varied by different contact ranges during cancerization (*Left*) and reversion (*Middle*), as well as their averages (*Right*).

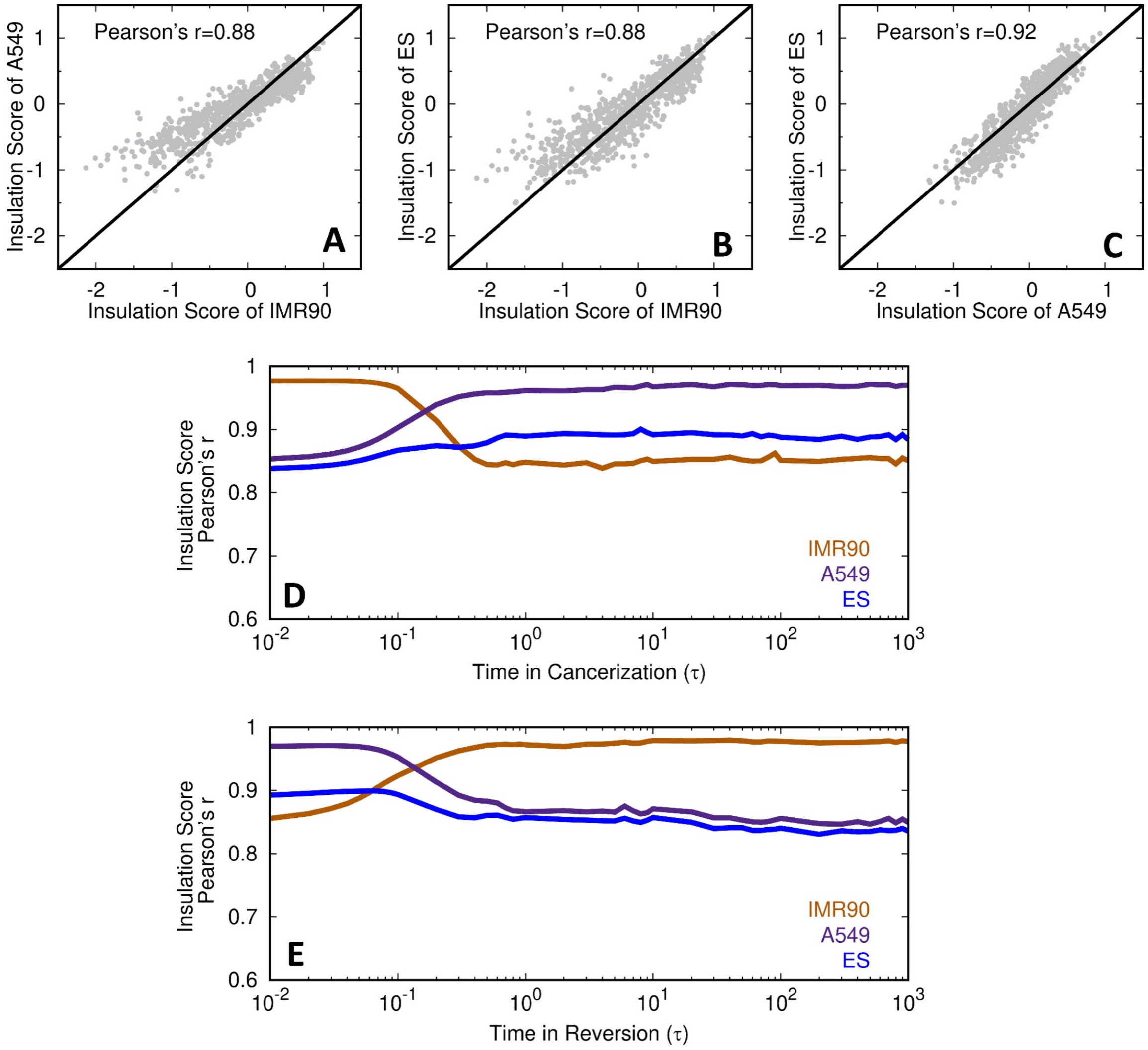


Figure S5: Insulation score. (A-C) The correlation between the insulation score among the IMR90, A549 and ES cell. The correlation coefficient of insulation score of the processing state during (D) cancerization and (E) reversion with that of the IMR90, A549 and ES cell.

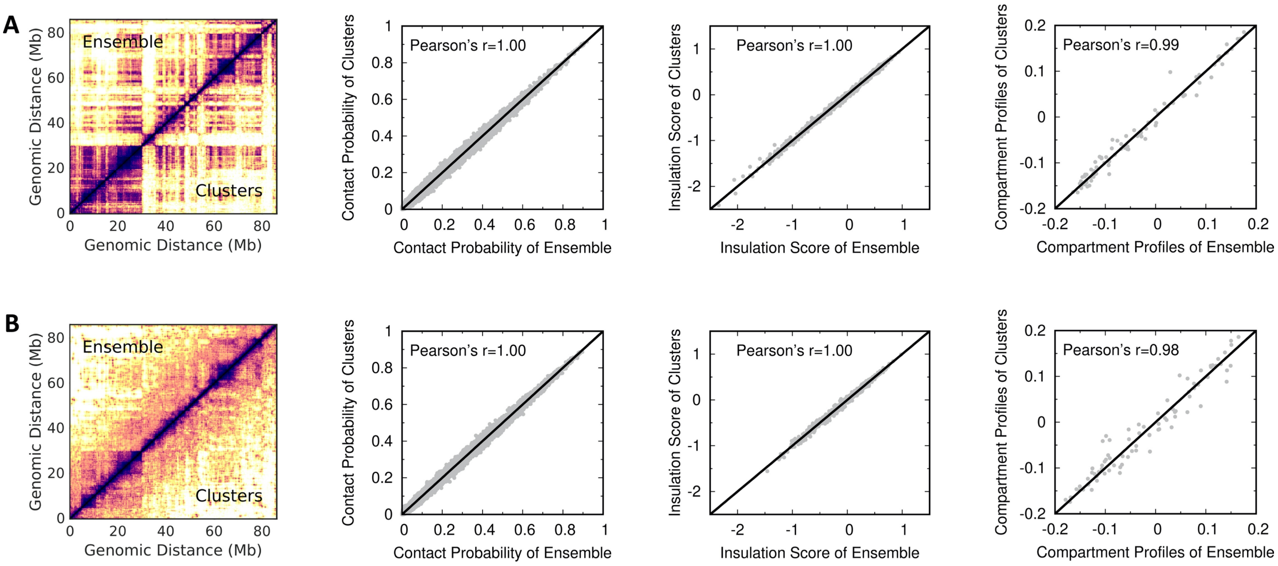


Figure S6: The comparisons of the chromosome structures between the clusters for initializing the landscape-switching model and the ensemble generated by the maximum entropy principle simulations. (A) The IMR90 cell. (B) The A549 cell.
